# Reengineering of an artificial protein cage for efficient packaging of active enzymes

**DOI:** 10.1101/2023.12.08.569934

**Authors:** Yusuke Azuma, Szymon Gaweł, Monika Pasternak, Olga Woźnicka, Elżbieta Pyza, Jonathan G. Heddle

**Author notes:** To whom correspondence should be addressed: Jonathan G. Heddle. These authors contributed equally to this work and are listed alphabetically (Szymon Gaweł, Monika Pasternak). Department of Biosciences, Durham University, South Road, DH1 3LE, United Kingdom (Jonathan G. Heddle).

## Abstract

Protein cages that readily encapsulate active enzymes of interest present useful nanotools for delivery and catalysis, wherein those with programmable disassembly characteristics serve as particularly attractive platforms. Here we establish a general guest packaging system based on an artificial protein cage, TRAP-cage, the disassembly of which can be induced by the addition of reducing agents. In this system, TRAP-cage with SpyCatcher moieties in the lumen was prepared using genetic modification of the protein building block and assembled into a cage structure with either monovalent gold ions or molecular crosslinkers. The resulting protein cage can efficiently capture guest proteins equipped with a SpyTag by simply mixing them in aqueous solution. This post-assembly loading system which circumvents the exposure of guests to thiol-reactive crosslinkers, enables packaging of enzymes possessing a catalytic cysteine or a metal cofactor while retaining their catalytic activity.

---

Encapsulation of active enzymes in a proteinaceous compartment is a powerful strategy for controlling their translocation in living systems as well as their catalytic activity. Nature has elegantly illustrated this concept, a primary example being viral capsids that co-package their genetic material with DNA/RNA-processing enzymes such as reverse transcriptase and deliver them together to target cells to initiate replication.^[1]^ Some bacteria localize a cluster of metabolic enzymes in protein shells, enhancing the pathway flux by sequestering intermediates and reducing undesired side reactions through selective internalization of target substrates, as represented by bacterial nano/microcompartments such as the carboxysome.^[2]^ These naturally occurring protein cages have inspired the design of smart nanocarriers and nanoreactors with prospective applications in molecular delivery and catalysis.^[3]^

Protein cages hold marked advantages as platforms for bioengineering.^[4]^ They include biocompatibility and monodispersity, as well as modifiability through chemical and genetic methods. Indeed, naturally occurring protein cages, such as virus-like particles (VLP), ferritin, and shell proteins of bacterial nano/microcompartments, have been extensively exploited as nanoscale reaction chambers for enzymes.^[3b,5]^ Proteinaceous compartments can also be constructed from protein building blocks that do not naturally form cage-like structures but gain such a capability upon appropriate engineering/modification.^[6]^ Such artificial protein cages have the potential to be endowed with structural and functional features that are unknown or even unfeasible in their natural equivalents.

We have established an artificial protein cage system based on the tryptophan RNA-binding attenuation protein, TRAP, a toroidal-shaped homo-undecamer protein derived from *Geobacillus stearothermophilus*.^[7]^ A TRAP variant possessing a cysteine at residue position 35, called TRAP^K35C^, can assemble with monovalent gold ions into a hollow cage-like structure composed of 24 copies of the ring-shaped subunits connected to each other via thiol-gold-thiol coordination (Figure. 1A).^[8]^ The resulting protein cage, referred to as TRAP^Au(I)^-cage, shows extremely high stability against heat and chaotropic reagents but can disassemble upon the addition of thiol- or phosphine-containing compounds.^[8a]^ This artificial protein cage, possessing triggerable disassembly characteristics, is a particularly attractive chamber for enzyme delivery,^[9]^ allowing the encapsulated guest molecules to be released at arbitrary timing and location, as required.

**Figure 1.**
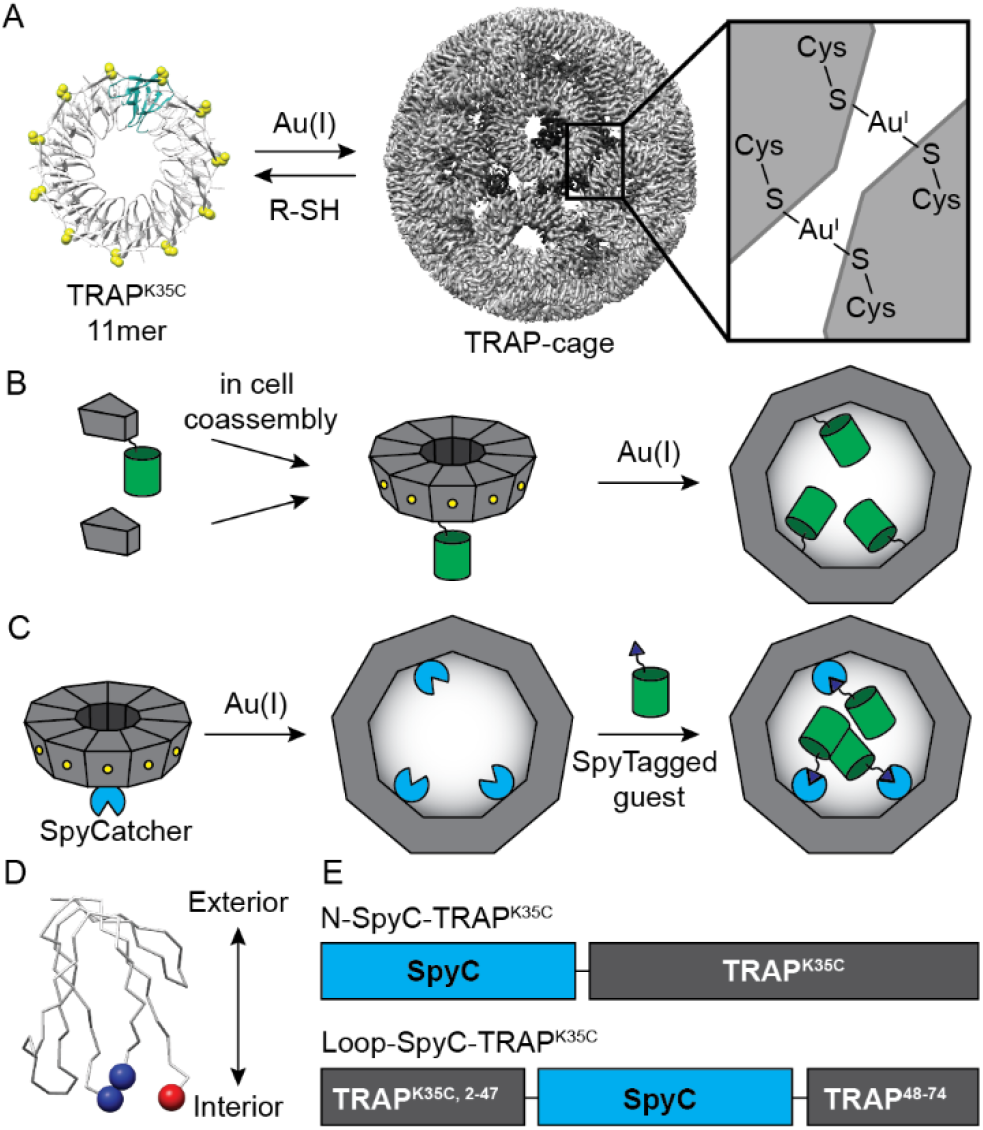
Post-assembly guest loading into TRAP-cages using SpyTag-SpyCatcher conjugation. (A) Au(I)-mediated assembly of TRAP^K35C^. The K35C mutation is shown as yellow spheres in the ribbon diagram of its 11-mer toroidal structure where a monomer unit is highlighted in cyan. (B) Guest packaging in TRAP^Au(I)^-cage using genetic fusion and patchwork formation. (C) Post-assembly guest loading into TRAP-cage using SpyTag-SpyCatcher conjugation. (D) Wire-diagram of TRAP monomer, where the positions of Thr3 (red) and Thr47/Glu48 (blue) are shown as spheres. (E) Design of TRAP^K35C^ variants containing SpyCatcher.

To package functional protein cargoes in the TRAP-cage lumen, we previously devised a strategy using genetic fusion and patchwork formation (Figure 1B).^[10]^ In this method, guest protein is genetically fused to the TRAP^K35C^ N-terminus which faces the cage interior when assembled^[8a,11]^ and coproduced with unmodified TRAP^K35C^ to form patchwork 11mer rings in cells.^[12]^ The resulting hetero TRAP-ring can be assembled with Au(I) into a cage-like structure containing the guest proteins in the lumen. However, the use of Au(I) to with the TRAP-guest fusion is likely to be incompatible with some enzymes, particularly those containing catalytic cysteine or metal cofactors.

To avoid guest exposure to gold ions, we modified the guest packaging system: protein building blocks are assembled first, followed by delivering guest proteins into the TRAP-cage lumen (Figure 1C). The TRAP^Au(I)^-cage wall has pseudosquare-shaped pores with side lengths of approx. 4 nm, formed by four undecameric building blocks. These are likely of sufficient size to allow entry of <4 nm proteins into the cage interior.^[8a]^ To capture target guest protein in the lumen, we employed the SpyTag-SpyCatcher system, a widely used bioconjugation method based on a split and engineered variant of a fibronectin-binding protein domain from *Streptococcus pyogenes*. The split fragments are able to form an intramolecular isopeptide bond spontaneously upon mixing in aqueous solution.^[13]^

The post-assembly loading system was first examined using TRAP^K35C^ fused to SpyCatcher at the TRAP N-terminus, N-SpyC-TRAP^K35C^, with a model cargo; a monomeric variant of green fluorescent protein (GFP) equipped with SpyTag at the GFP N-terminus, SpyT-msfGFP.^[14]^ The N-SpyC-TRAP^K35C^ variant was coproduced with untagged TRAP^K35C^ to form a patchwork ring structure in host *Escherichia coli* cells, followed by cage assembly with Au(I) as previously reported (Figure 1D,E and Figure S1A).^[10]^ The obtained TRAP-cage, referred to as N-SpyC-TRAP^Au(I)^-cage, was then mixed in phosphate-buffered saline (PBS) with SpyT-msfGFP at a SpyTag:SpyCatcher ratio of 1:1. Size-exclusion chromatography of the mixture showed a peak corresponding to the assembled TRAP-cage with substantial absorbance at 488 nm, suggesting successful complex formation (Figure S2A). Covalent bond formation between N-SpyC-TRAP^K35C^ and SpyT-msfGFP was confirmed by sodium dodecyl sulfate-polyacrylamide gel electrophoresis (SDS-PAGE) of the isolated particles (Figure 2A). However, native-PAGE showed a clear band shift upon conjugation, suggesting a change in the size of the cage assemblies (Figure 2B). Negative-stain transmission electron microscopy (TEM) confirmed the exterior surface became rougher compared to the sample before the addition of SpyT-msfGFP (Figure 2C). These results suggested that the guests are partially displayed on the TRAP-cage exterior, instead of or in addition to encapsulation in the lumen.

**Figure 2.**
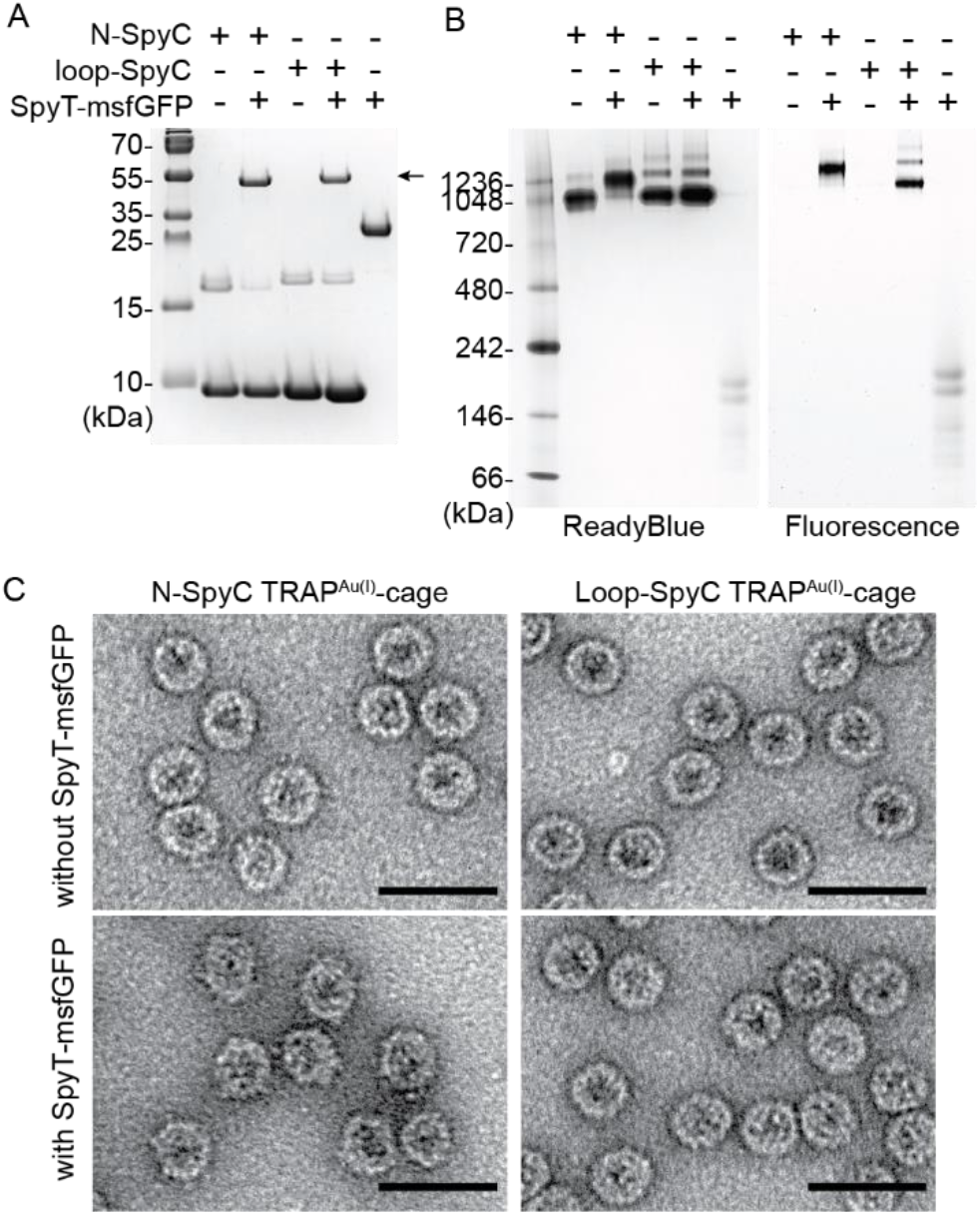
GFP encapsulation. (A-C) SDS-PAGE (A), Native-PAGE (B), and TEM (C) analysis of filled TRAP-cages. The theoretical molecular masses of each protein are as follows: TRAP^K35C^, 8.5 kDa; SpyC-TRAP^K35C^(N-SpyC), 18.4 kDa; TRAP^K35C^-loopSpyC (loopSpyC), 19.1 kDa; SpyT-msfGFP, 28.7 kDa. Arrow indicates the bands corresponding to the TRAP-msfGFP conjugates (47.8 kDa). Scale bar = 50 nm.

We assumed that this potential guest leakage could be caused by SpyCatcher entering the central pore of the TRAP 11mer ring (Figure S3). The pore diameter is ∼2 nm, similar in size to SpyCatcher (Figure S3A).^[11,15]^ Additionally, the TRAP N-terminus locates at the pore region. This probably allows transient external localization of the SpyCatcher moiety, allowing an opportunity for conjugation with SpyT-msfGFP and subsequent (partial) externalization (Figure S3B).

To remove the chance of external display of the guest protein, the SpyCatcher moiety was next introduced between residues Thr47 and Glu48 of TRAP^K35C^, a loop region facing the interior of the cage assembly (Figure 1D,E).^[8a,11]^ Despite this substantial addition (92 amino-acid-length SpyCatcher is inserted to the middle of TRAP monomer sequence being only 74 amino acids), the resulting variant, called Loop-SpyC-TRAP^K35C^, successfully expressed in *E. coli* cells to form patchwork structures with TRAP^K35C^, and assembled with Au(I) into a cage-like structure, referred to as Loop-SpyC-TRAP^Au(I)^-cage (Figure S1B). Guest packaging using this TRAP-cage was tested as for the previous cage, and showed successful encapsulation in the lumen with no evidence of leakage at the exterior. The Loop-SpyC-TRAP^Au(I)^-cages are able to form a complex with SpyT-msfGFP via a covalent bond without any notable morphology changes (Figure 2 and Figure S2B).

The number of GFP associated with TRAP-cages was quantified using the absorbance ratio at 280/488 nm,^[5d]^ giving 31 ± 7 and 18 ± 3 guests per cage for N-SpyC-TRAP^Au(I)^- and Loop-SpyC-TRAP^Au(I)-^cages, respectively (Figure S4A,B). These numbers correspond to 71 ± 7 % and 94 ± 2% of SpyCatcher moieties reacting with SpyT-msfGFP (Figure S4C). The high modification efficiency observed for N-SpyC-TRAP^Au(I)^-cage likely reflects the fact that the SpyCatcher moiety can be displayed on the cage exterior, giving almost no spatial restrictions for the reaction. In contrast, ∼18 GFP likely represents the maximum loading capacity of Loop-SpyC-TRAP^Au(I)^-cage due to the limited space to accommodate the guests. This was supported by experiments in which the guest ratio was increased to 1.5 equivalent (20 μM SpyCatcher and 30 μM SpyT-msfGFP) and which did not result in a higher encapsulation efficiency, while the loading is nearly quantitative at lower guest stoichiometry (5 or 10 μM) (Figure S5A).

Temperature plays a role in the maximum loading of Loop-SpyC-TRAP^Au(I)^-cage. When the host and guest were mixed in 1:1 ratio at 4 °C, the reaction reached a plateau of ∼50% efficiency in 4 hours, defined as conjugated SpyC per total SpyC estimated by SDS-PAGE densitometry assay (Figure S5B, blue). A similar reaction rate was observed at room temperature. However, extension of the reaction to 20 hours drove the reaction efficiency up to ∼70% (Figure S5B, black). Thermal motion of proteins may contribute to promoting a higher guest density in the TRAP-cage lumen.

Having successfully encapsulated GFP in TRAP-cage, we next tested if the system is applicable to packaging of active enzymes possessing a catalytic cysteine. As a model, an engineered variant of sequence-specific protease derived from tobacco etch virus, sTEVp,^[16]^ was employed. As for GFP, this enzyme was equipped with Spy-tag at the N-terminus, SpyT-sTEVp, and mixed in PBS with Loop-SpyC-TRAP^Au(I)^-cages. Successful encapsulation was confirmed by native-PAGE, SDS-PAGE, and TEM (Figure S6). However, upon encapsulation in the TRAP^Au(I)^-cages, the enzyme showed no detectable cleavage activity against a peptide containing the recognition sequence (Figure S7, S8).^[17]^ Notably, the guest enzyme became active again immediately after the addition of dithiothreitol (DTT) (Figure S8, red). A similar enzyme inactivation and recovery by DTT was also observed when the free enzyme was treated with Au(I) (Figure S8, black). Based on these results, we assumed that the guest enzyme may strip some bridging gold ions from the TRAP-cage wall via coordination with the catalytic cysteine.

To verify the Au(I)-mediated quenching hypothesis as well as to achieve encapsulation of active enzyme, we switched the TRAP-cage assembly system from Au(I) to dithiobismaleimidoethane (DTME) (Figure S9).^[10]^ We previously reported that the bismaleimide molecular crosslinker can induce TRAP-cage assembly, referred to as TRAP^DTME^-cage, in a similar manner as Au(I) but via Michael addition instead of metal coordination. Since DTME contains a disulfide bond in the structure, the resulting cage can be disassembled into fragments by the addition of reducing agents. When SpyT-sTEVp was loaded into TRAP^DTME^-cage possessing SpyCatcher moieties, Loop-SpyC-TRAP^DTME^-cage (Figure 3A,B and Figure S10A), the guest enzyme exhibited substantial activity as expected (Figure 3C).

**Figure 3.**
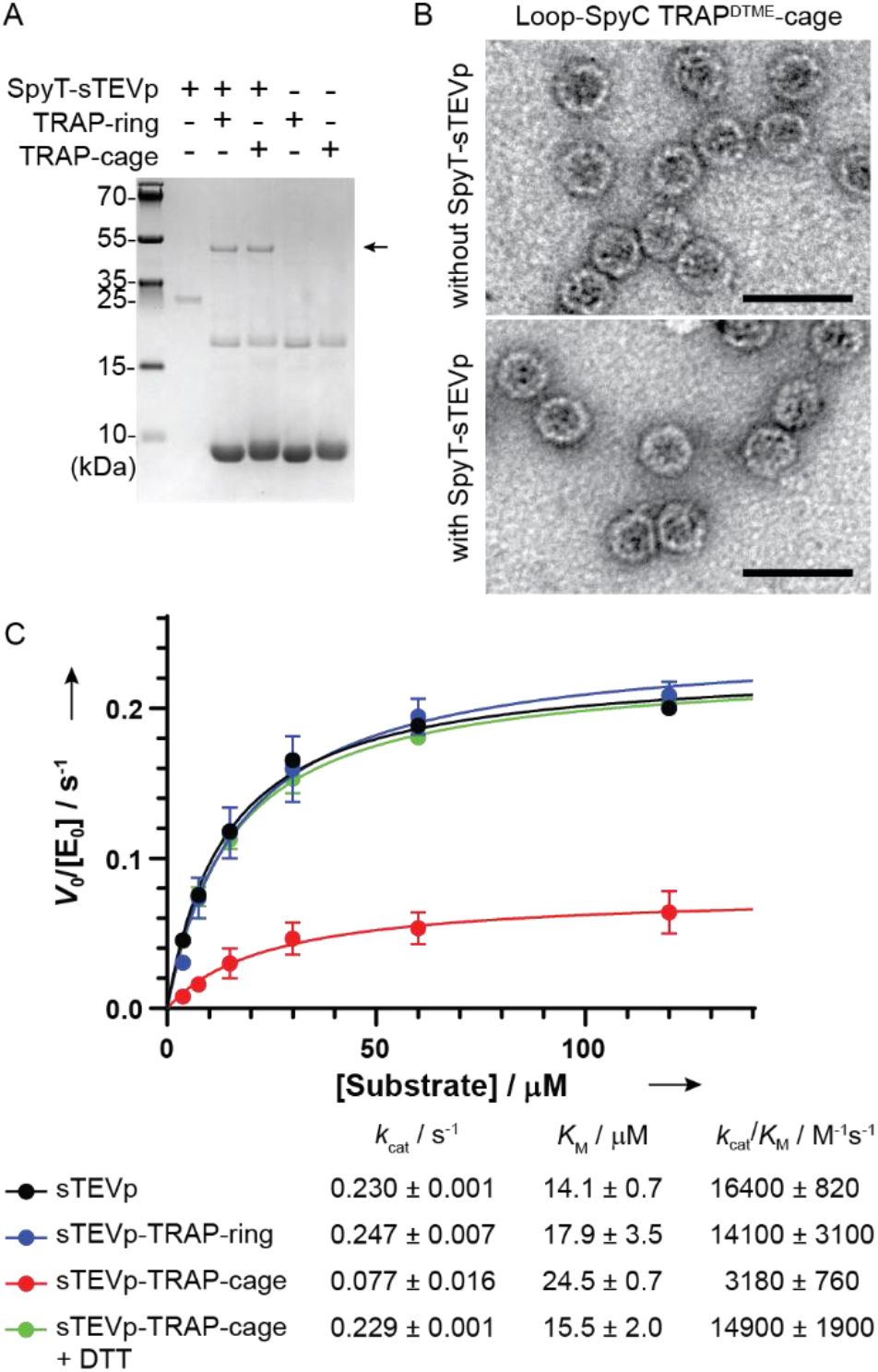
TEV protease encapsulation. (A,B) SDS-PAGE (A) and negative-stain TEM (B) analysis of Loop-SpyC-TRAP^K35C^ conjugated with SpyT-sTEVp in the form of 11mer ring (TRAP-ring) or DTME-mediated cage assembly (TRAP-cage). The theoretical molecular masses of each protein are as follows: TRAP^K35C^, 8.5 kDa; Loop-SpyC-TRAP^K35C^, 19.1 kDa; SpyT-sTEVp, 30.6 kDa. Arrow indicates the bands corresponding to the TRAP-sTEVp conjugates (49.7 kDa). Scale bar = 50 nm. (C) Michealis-Mentan plots for the activity of SpyT-sTEVp. Data are presented as means ± standard deviations from two independent experiments.

Encapsulation of sTEVp in Loop-SpyC-TRAP^DTME^-cages altered the apparent steady-state kinetics parameters for the proteolytic reaction. Compared to those of SpyT-sTEVp free in solution, a ∼3-fold decrease in *k*_cat_ and a 1.7-fold increase in *K*_M_ were observed (Figure 3C, black and red). Conjugation to TRAP-ring via SpyTag-SpyCatcher is not the reason for the change as proven by the fact that conjugation of the SpyT-sTEVp with Loop-SpyC-TRAP^K35C^-containing 11mer ring had almost no effect on the enzyme activity (Figure 3C, black and blue). Notably, a similar decrease in TEV protease activity has been shown when the enzyme was encapsulated in another protein cage formed by an engineered variant of lumazine synthase with a local guest enzyme concentration of ∼8 mM.^[17]^ Although we kept the average guest number per cage relatively low (6 enzymes per cage), the local concentration still reached ∼3 mM assuming the interior volume of TRAP^DTME^-cage is ∼3000 nm^3^.^[10]^ The change in the kinetic parameters is likely attributable to the high enzyme density within the cages resulting in partial aggregation of the guest enzyme.

sTEVp inactivation upon encapsulation in TRAP-cages is reversible. This was tested by exploiting the fact that TRAP^DTME^-cages can be opened: sTEVp-TRAP-cages were treated with DTT after encapsulation. Successful disassembly of the TRAP-cages was confirmed by Native-PAGE and TEM imaging (Figure S10). This sTEVp release from the cage assembly resulted in close to 100% recovery of the guest enzyme activity (Figure 3C, green).

Influence of encapsulation in TRAP^DTME^-cages on enzyme activity appears to depend on the guests. In addition to sTEVp, we tested with another cysteine protease/ligase, an engineered variant of sortase A, SrtA,^[18]^ as well as a metalloenzyme, human carbonic anhydrase II, hCAII.^[19]^ Upon encapsulation in Loop-SpyC-TRAP^DTME^-cages (Figure S11), these enzymes retained a catalytic efficiency close to that of the controls, consisting of the corresponding enzymes free in solution (Figure S12).

In summary, this study establishes that TRAP-cage possessing recombinantly inserted SpyCatcher moieties in an interior-facing loop region can function as a general and robust platform for packaging functional proteins equipped with a SpyTag. The loading procedure is easy and efficient. Simple mixing of the host cage and guest in an aqueous solution allows the inclusion complex formation in a nearly quantitative manner, up to ∼18 guests per cage in the case of GFP. TRAP-cage formed using an Au(I)-mediated assembly system was found to be unsuitable for packaging enzymes possessing a catalytic cysteine which potentially removes metal ions from the protein cage wall resulting in self-inactivation. This issue can be overcome using bismaleimide as an alternative crosslinker. Such protein cages containing active enzymes may serve as smart nanoreactors that sort substrates based on their size and/or chemical properties.^[17,20]^ In this regard, triggerable disassembly of the TRAP^DTME^-cage offers a practical and convenient approach for investigating the effect of confinement on catalytic activity.^[3b,5c]^ Moreover, TRAP-cages, being able to readily package and release guest proteins, may provide an ideal in vivo tissue-specific delivery vehicle upon further decoration of the exterior using chemical or genetic methods.^[9]^ The sequence-specific proteases used in this study could be replaced with examples with the potential to control both endogenous, e.g. caspase,^[21]^ or artificial, cellular signaling events.^[22]^ The possibility of using modular protein cages for the intracellular delivery of active enzymes is currently under investigation in our laboratory.

## Supporting Information

Size exclusion chromatograms, schematic representations of guest leakage mechanism and cage formation via molecular crosslinkers, guest packaging quantification and kinetics, substrate structures, electrophoresis and TEM images, enzyme kinetics, and detailed descriptions for the materials and methods are available in supporting information.^[23-28]^

## Acknowledgements

This work was generously supported by Jagiellonian University and the National Science Centre (NCN) Poland (Symfonia 2016/20/W/NZ1/00095, Maestro 2019/34/A/NZ1/00196, and Preludium 2021/41/N/NZ1/02635).

## Conflict of interest

The authors declare the following competing interests: The authors are named on the following patents: WO 2020/035716 A1 (J.G.H.), WO 2022/182260 A1 (J.G.H., Y.A. and S.G.), WO 2022/182261 A1 (J.G.H. and Y.A.), WO 2022/182262 A1 (J.G.H. and Y.A.). J.G.H. is also the founder of and holds equity in nCage Therapeutics LLC, which aims to commercialize protein cages for therapeutic applications. Y.A. is also a member of the scientific advisory board of nCage Therapeutics. S.G. is an employee of nCage Therapeutics. The authors declare that they have no other competing interests.

