## Supporting Information for "Reengineering of an artificial protein cage for efficient packaging of active enzymes"

**Index**

**1. Supporting data**

**Figure S1. Purification of Au(I)-mediated assembly of TRAP$^{K35C}$ containing SpyCatcher. ---- S3**

**Figure S2. Purification of TRAP-cages containing GFP. --------------------------------------------- S4**

**Figure S3. Potential mechanism of the guest display on the TRAP-cage exterior. --------------- S5**

**Figure S4. Quantification of GFP encapsulated in TRAP-cages. ------------------------------------ S6**

**Figure S5. Stoichiometry control and reaction kinetics of GFP packaging. ----------------------- S7**

**Figure S6. TEV protease encapsulation in TRAP$^{Au(I)}$-cage. ------------------------------------------ S8**

**Figure S7. Substrates used for enzyme kinetics. -------------------------------------------------------- S9**

**Figure S8. Inactivation of TEV protease by TRAP$^{Au(I)}$-cage and Au(I). --------------------------- S10**

**Figure S9. DTME-mediated TRAP-cage formation. ------------------------------------------------ S11**

**Figure S10. TEV protease encapsulation in TRAP$^{DTME}$-cage and its release by DTT. ---------- S12**

**Figure S11. SrtA and hCAII encapsulation in TRAP$^{DTME}$-cage. ------------------------------------ S13**

**Figure S12. SrtA and hCAII activity in TRAP$^{DTME}$-cage. ------------------------------------------- S14**

**2. Materials and methods**

**Materials ---------------------------------------------------------------------------------------------------- S15**

**Molecular cloning ------------------------------------------------------------------------------------------ S16**

**Table S3. Oligonucleotides used in this study ---------------------------------------------- S18**

**Table S4. Plasmids used for protein production in this study ---------------------------- S19**

**Table S5. Proteins used in this study ---------------------------------------------------------- S20**

**Protein production and purification. ---------------------------------------------------------------- S22**

**TRAP-cage assembly ------------------------------------------------------------------------------------- S23**

**GFP encapsulation in TRAP$^{Au(I)}$-cage ----------------------------------------------------------------- S24**

**Negative-stain transmission electron microscopy (TEM) ------------------------------------------ S24**

**Native-PAGE ----------------------------------------------------------------------------------------------- S24**

**SDS-PAGE -------------------------------------------------------------------------------------------------- S25**

**GFP encapsulation efficiency --------------------------------------------------------------------------- S25**

**Enzyme encapsulation ----------------------------------------------------------------------------------- S25**

**Enzyme kinetics ------------------------------------------------------------------------------------------- S26**

**3. References ---------------------------------------------------------------------------------------------- S27**

### 1. Supporting data

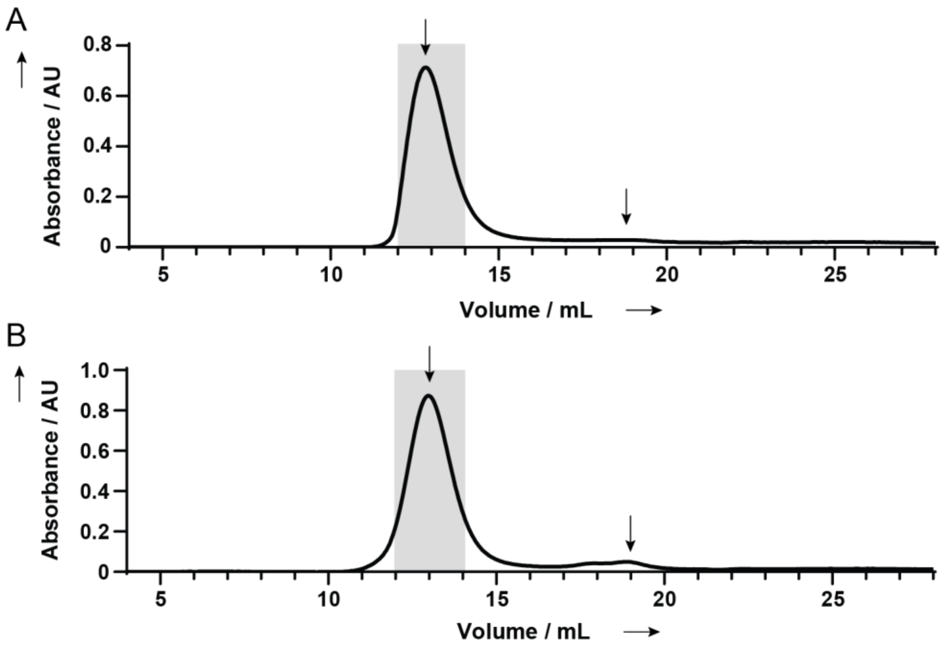


**Figure S1. Purification of Au(I)-mediated assembly of TRAP$^{K35C}$ containing SpyCatcher.** (A,B) Size-exclusion chromatogram of patchwork undecamer ring composed of TRAP$^{K35C}$ and N-SpyC-TRAP$^{K35C}$ (A) or Loop-SpyC-TRAP$^{K35C}$ (B) after overnight incubation with TPPMS-Au(I)-Cl (1 equiv.). The fractions used for subsequent characterization are marked grey. The peak at ~13 and 19 mL, indicated by arrows, corresponds to cage-like structure and undecamer ring, respectively.

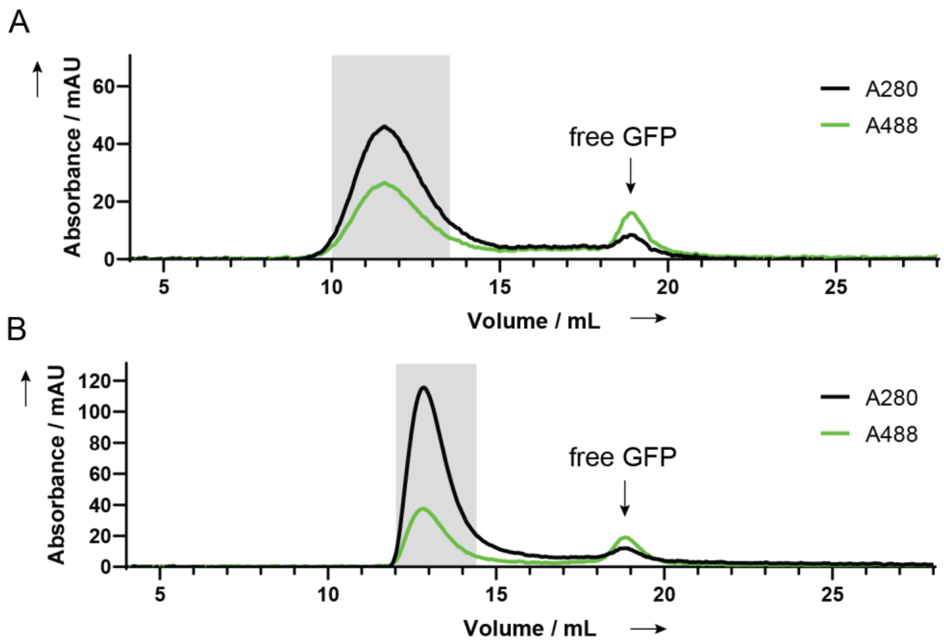


**Figure S2. Purification of TRAP-cages containing GFP.** (A,B) Size-exclusion chromatogram of N-SpyC-TRAP$^{Au(I)}$-cages (A) and LoopSpyC-TRAP$^{Au(I)}$-cages (B) (both 20 µM with respect to SpyCatcher concentration) after overnight incubation with SpyT-msfGFP (20 µM). The fractions used for subsequent characterization are marked grey. The peak at ~19 mL corresponds to free SpyT-msfGFP that remained unreacted.

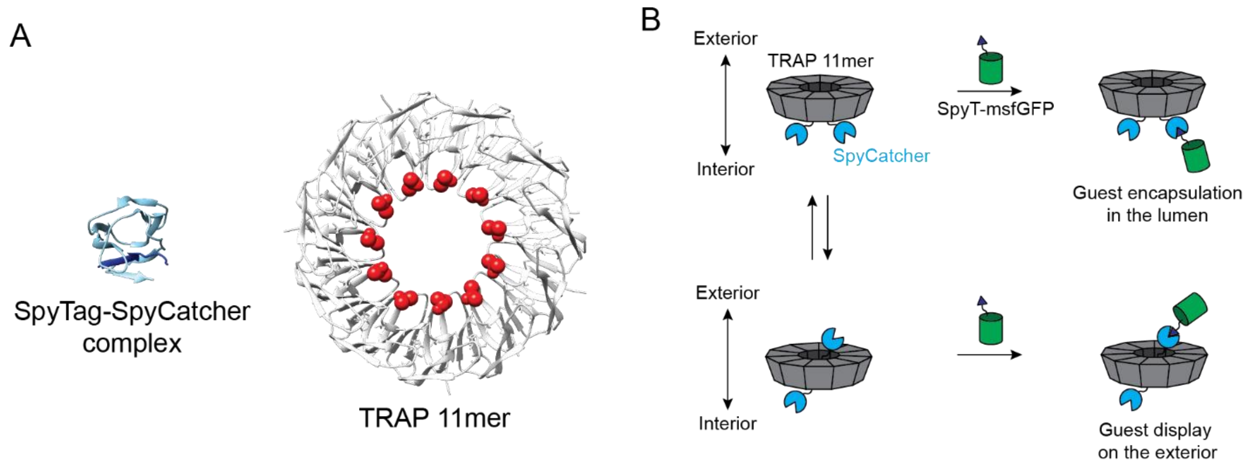


**Figure S3. Potential mechanism of the guest display on the exterior of TRAP-cage.** (A) Structural comparison of SpyTag-SpyCatcher complex (PDB: 4MLI) and TRAP 11mer (1UTD). The threonine residue at position 3 is highlighted as red spheres. (B) Hypothetical mechanism of SpyCatcher translocation through the central pore of TRAP 11mer ring, determining the position of guest GFP after SpyTag-SpyCatcher conjugation.

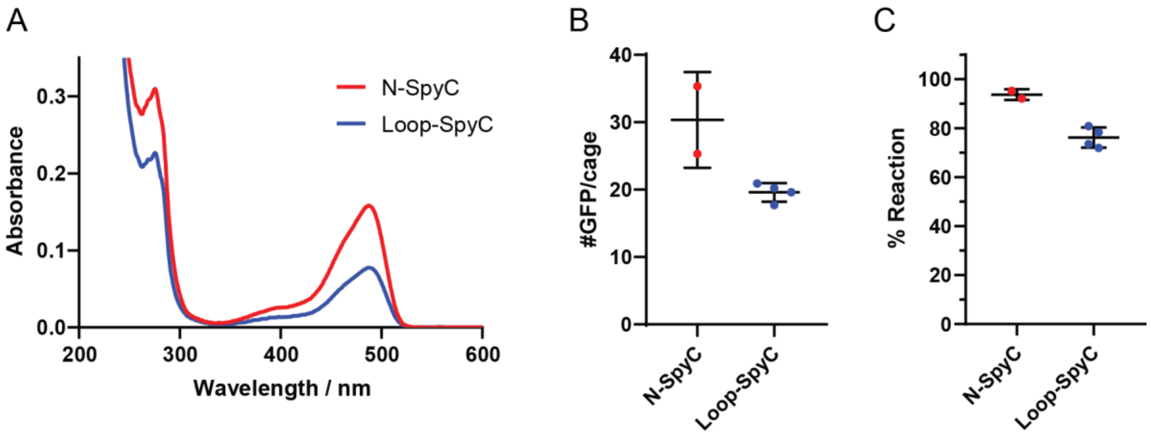


**Figure S4. Quantification of GFP encapsulated in TRAP-cages.** (A) Absorbance spectra of N-SpyC-TRAP$^{Au(I)}$-cages (N-SpyC) and LoopSpyC-TRAP$^{Au(I)}$-cages (Loop-SpyC) containing SpyT-msfGFP. (B) The number of GFP per cage was estimated from the absorbance ratio at 280 and 488 nm. (C) Encapsulation efficiency. Percent reaction is defined as (number of GFP per cage)/(number of SpyC per cage)×100, where the number of GFP and SpyC was determined by absorbance spectra and SDS-PAGE densitometry assay, respectively.

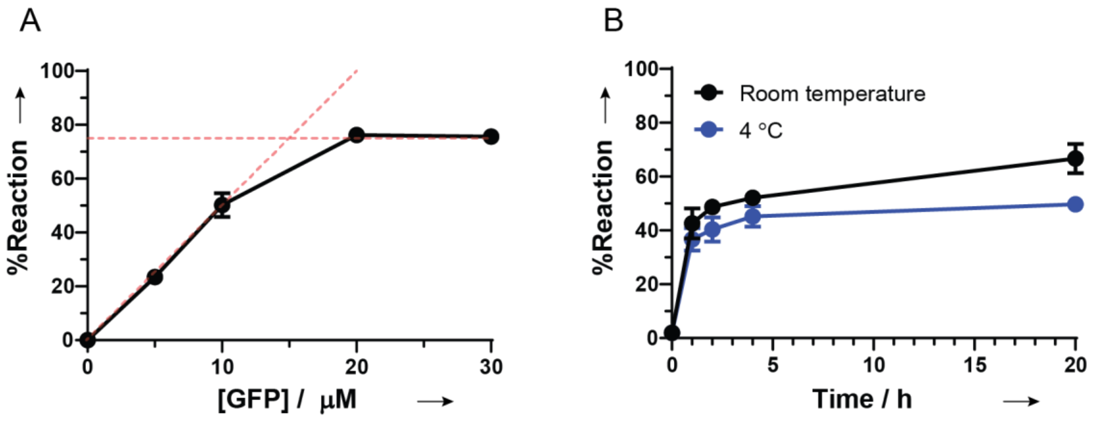


**Figure S5. Stoichiometry control and reaction kinetics of GFP packaging.** (A) Titration of SpyT-msfGFP (0, 5, 10, 20, and 30 μM) onto Loop-SpyC-TRAP$^{Au(I)}$-cages (20 μM with respect to SpyCatcher concentration). Percent reaction is defined as (number of SpyCatcher conjugated with SpyT-msfGFP)/(number of total SpyCatcher)×100, estimated by SDS-PAGE densitometry assays. Red dashed lines indicate a trend line expected for the case if 100% input SpyT-msfGFP is conjugated as well as the saturation point in % reaction. (B) Monitoring the conjugation reaction using SpyT-msfGFP (20 μM) and Loop-SpyC-TRAP$^{Au(I)}$-cages (20 μM with respect to SpyCatcher concentration) at 0, 1, 2, 4, and 20 hours at either room temperature or 4 °C.

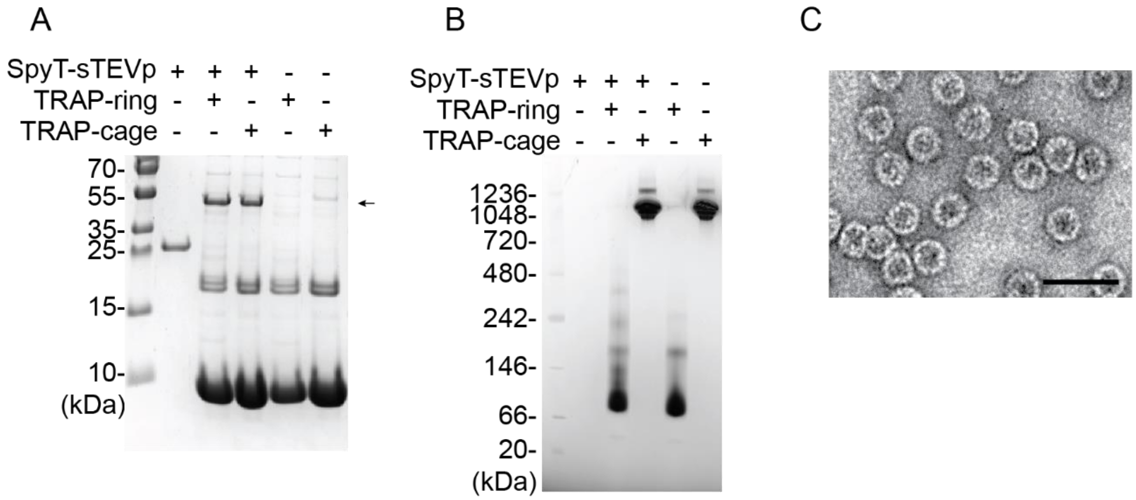


**Figure S6. TEV protease encapsulation in TRAP$^{Au(I)}$-cage.** (A) SDS-PAGE (B), Native-PAGE, and (C) TEM analysis of filled TRAP-cages. The theoretical molecular masses of each protein are as follows: TRAP$^{K35C}$, 8.5 kDa; Loop-SpyC-TRAP$^{K35C}$, 19.1 kDa; SpyT-sTEVp, 30.6 kDa. Arrow indicates the bands corresponding to TRAP-sTEVp conjugates (49.7 kDa). Scale bar = 50 nm.

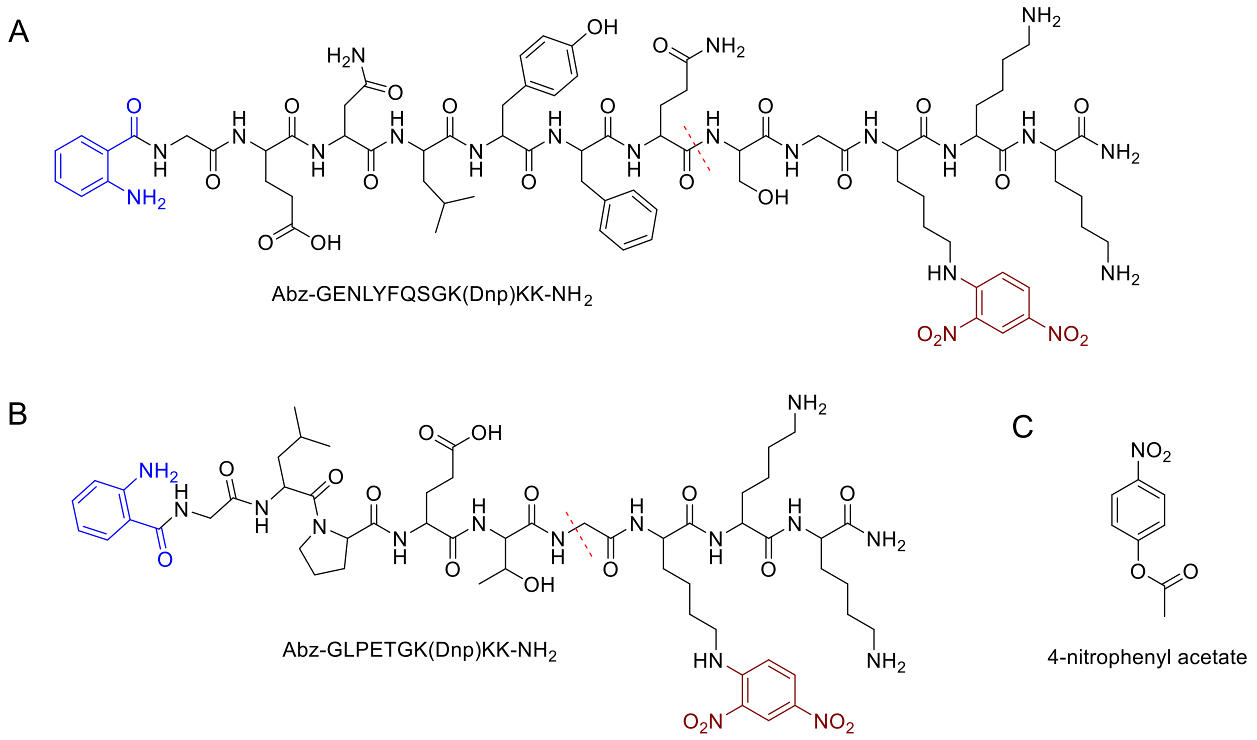


**Figure S7. Substrates used for enzyme kinetics.** (A,B) The substrate peptides containing N-terminal aminobenzoic acid (Abz, shown in blue) and C-terminal 2,4-dinitrophenyl (Dnp, shown in brown) lysine across the recognition sequence of TEV protease (A) and sortase A (B). These peptides possess two lysine residues at the C-termini to increase solubility in water. The cleavage site is indicated with red dashed lines. Cancellation of Förster resonance energy transfer (FRET) upon cleavage increases the fluorescent emission of Abz, which is used for monitoring the activity of the peptidases. (C) A chromogenic esterase substrate. The absorbance change upon hydrolysis to yield 4-nitrophenol was used for monitoring carbonic anhydrase activity in this study.

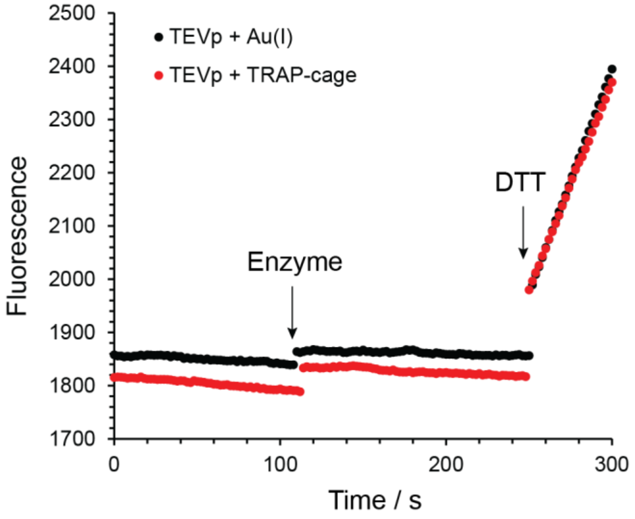


**Figure S8. Inactivation of TEV protease by TRAP$^{Au(I)}$-cage and Au(I).** The sTEVp (8 µM) was mixed with either Loop-SpyC-TRAP$^{Au(I)}$-cage (36 µM with respect to SpyCatcher concentration) or TPPMS-Au(I)-Cl (80 µM), and then kept at 4 °C overnight. The substrate peptide (30 µM) was added with the enzyme (final 40 nM) and then DTT (final 5 mM). An increase in fluorescent signal attributed to the FRET peptide cleavage was observed only after the addition of DTT.

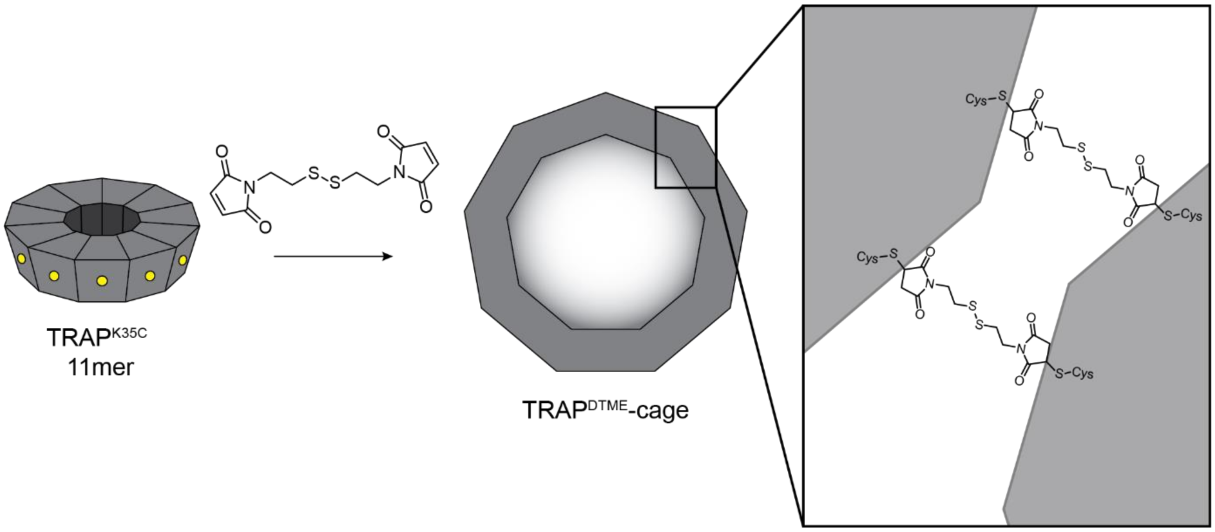


**Figure S9. DTME-mediated TRAP-cage formation.** TRAP$^{K35C}$ 11mer ring can assemble into a cage-like structure through reaction with bismaleimide crosslinkers such as DTME. Images are not scaled.

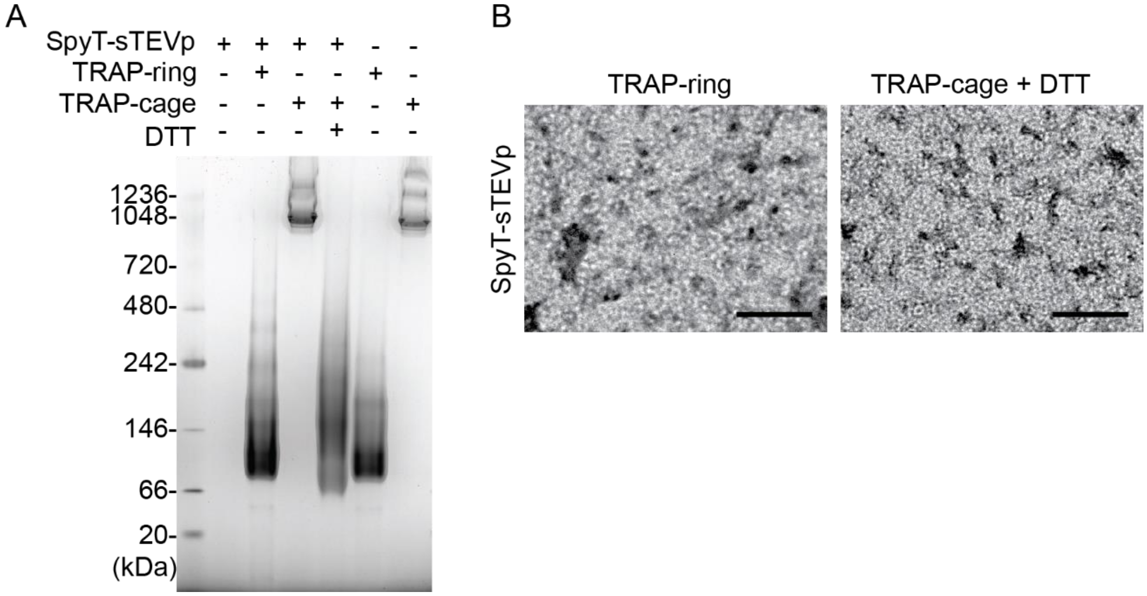


**Figure S10. TEV protease encapsulation in TRAP$^{DTME}$-cage and its release by DTT.** (A) Native-PAGE analysis of TRAP$^{K35C}$-loopSpyC conjugated with SpyT-sTEVp in the form of 11mer ring (TRAP-ring) or cage assembled with DTME (TRAP-cage). The enzyme alone did not migrate into the gel, likely due to the positively charged surface (*p*I = 10.3). (B) Negative-stain TEM images of TRAP$^{K35C}$-loopSpyC conjugated with SpyT-sTEVp in the form of 11mer ring (TRAP-ring) or cage assembled with DTME followed by treatment with 10 mM DTT overnight (TRAP-cage + DTT). Scale bar = 50 nm.

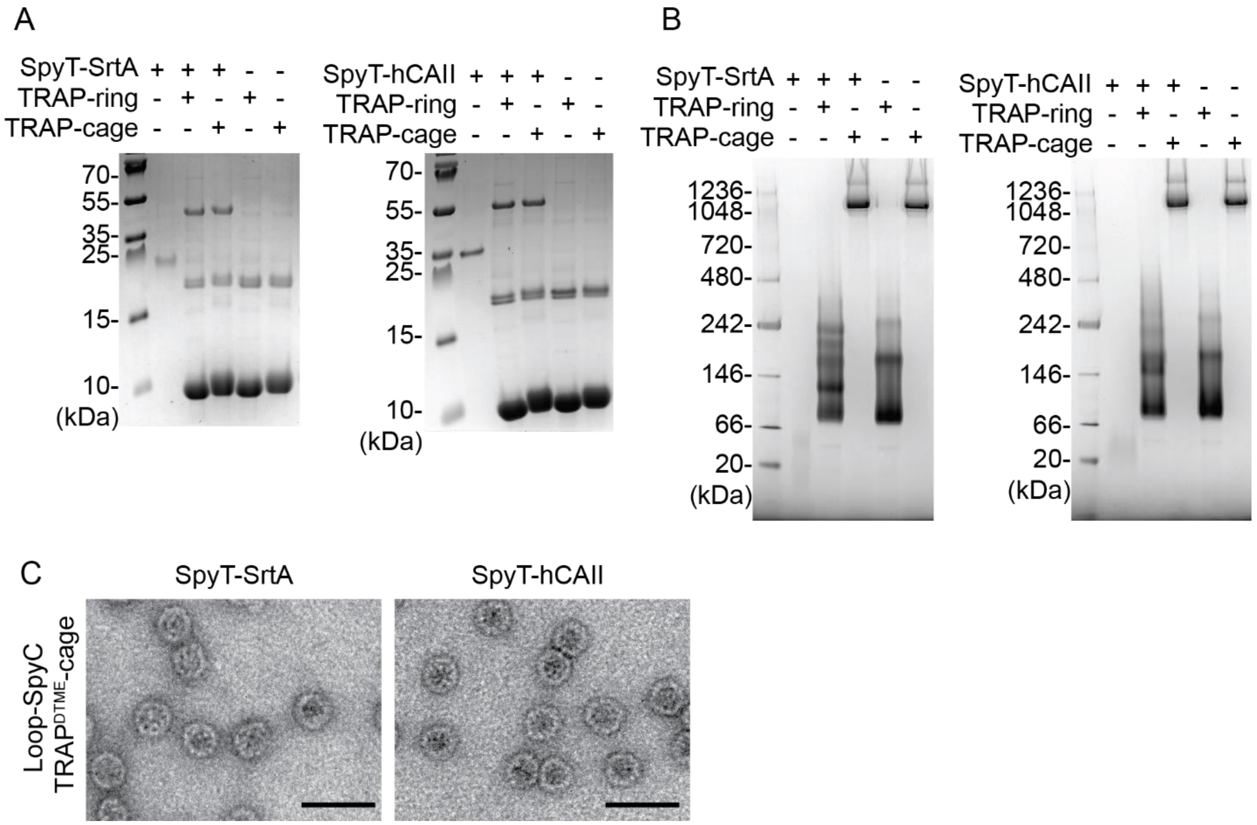


**Figure S11. SrtA and hCAII encapsulation in TRAP$^{DTME}$-cage.** (A) SDS-PAGE, (B) Native-PAGE, and (C) negative-stain TEM analysis of Loop-SpyC-TRAP$^{K35C}$ conjugated with SpyT-SrtA or SpyT-hCAII in the form of 11mer ring (TRAP-ring) or cage assembled with DTME (TRAP-cage). The theoretical molecular masses of each protein are as follows: TRAP$^{K35C}$, 8.5 kDa; Loop-SpyC-TRAP$^{K35C}$, 19.1 kDa; SpyT-SrtA, 20.2 kDa; SpyT-hCAII, 32.3 kDa; Loop-SpyC-TRAP$^{K35C}$-SpyT-SrtA, 39.3 kDa; Loop-SpyC-TRAP$^{K35C}$-SpyT-hCAII, 51.4 kDa. Scale bar = 50 nm.

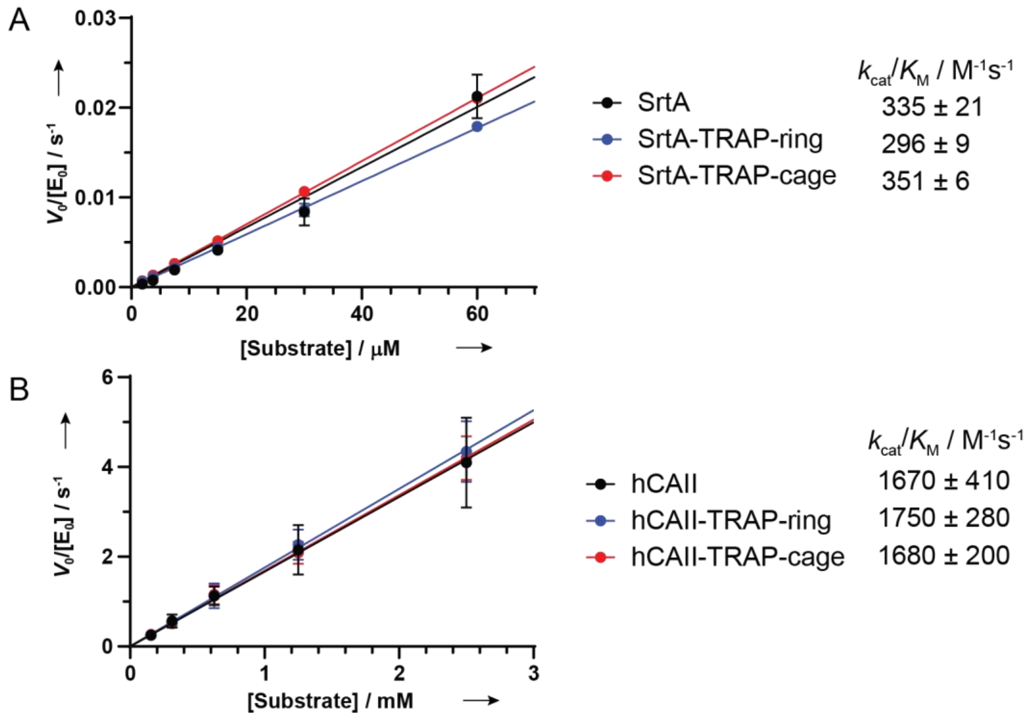


**Figure S12. SrtA and hCAII activity in TRAP$^{DTME}$-cage.** The kinetic efficiency, $k_{cat}/K_M$, of these enzymes was determined by linear fitting of the data points at the substrate concentration no greater than 60 µM or 2.5 mM for SrtA and hCAII, respectively. Data are presented as means ± standard deviations from two independent experiments.

### 2. Materials and methods

#### Materials

Chloro[diphenyl(3-sulfonatophenyl)phosphine]gold(I) sodium salt hydrate was purchased from Strem Chemicals UK Ltd. (Cambridge, UK) and reconstituted in 50 mM sodium phosphate buffer (pH 8.0) to 5 mM stock concentration before use. Dithiobismaleimidoethane (DTME), Phusion High-Fidelity DNA Polymerase, GeneJET Plasmid Miniprep Kit, GeneJET Gel Extraction Kit, and HisPur Ni-NTA resin were purchased from Thermo Fisher Scientific (Waltham, MA, USA). The molecular cross-linker was reconstituted in dimethyl formamide (DMF) to 50 mM stock concentration prior to use. Restriction enzymes and T4 DNA ligase used for molecular cloning were purchased from New England BioLabs (Ipswich, MA, USA). Isopropyl-β-D-thiogalactopyranoside (IPTG), Kanamycin sulfate, and dithiothreitol (DTT) were purchased from VWR (Radnor, PA, USA). cOmplete mini EDTA-free protease-inhibitor cocktail was from Roche (Basel, Switzerland). Ampicillin sodium salt, urea, 20% sodium dodecyl sulfate (SDS) solution were purchased from Lab Empire (Rzeszów, Poland). Chloramphenicol was purchased from PanReac AppliChem (Darmstadt, Germany). Ethylenediaminetetraacetic acid (EDTA) disodium salt dihydrate, L-tryptophan, lysosyme, deoxyribonuclease I from bovine pancreas (DNase I), tetracycline, 4-nitrophenyl acetate, and triglycine (Gly-Gly-Gly) were purchased from Sigma Aldrich (St. Louis, MO, USA). The plasmids, pACYC_H-sfGFP, pBAD_H-tev-GFP(+19), and pACYC_H-GFP(-30), were kind gifts from Prof. Donald Hilvert at ETH Zurich, Switzerland. Bacterial expression vectors for TRAP containing the K35C mutation (pET21_TRAP-K35C), SpyCatcher (pET28_H-SpyCat), super TEV protease (pMAL_H-SpyT-sTEVp-R5),[16] an engineered sortase A (pET30_H-SrtA*),[18] and human carbonic anhydrase II (pET28_H-SpyT-CAII) were prepared by BioCat GmbH (Heidelberg, Germany). All the oligonucleotides were synthesized by Sigma Aldrich. The substrate peptides for protease assays were synthesized and characterized using reverse-phase high-performance liquid chromatography (RP-HPLC) and electron spray ionization mass spectrometry (ESI-MS) by GenScript Biotech (Piscataway, NJ, USA): Abz-GENLYFQSGK(Dnp)KK-$NH_2$, observed molecular mass 1682.4 (1682.8, theoretical) and purity 96.3%; Abz-GLPETGK(Dnp)KK-$NH_2$, observed molecular mass 1241.4 (1241.4, theoretical) and purity 98.7%.

### Molecular cloning

The plasmids, oligonucleotides, and protein sequences used in this study are summarized in Table S1-3. *E. coli* strain NEB5α was used as the host cell for every cloning step. Sequences of plasmids were confirmed by DNA Sanger sequencing performed by Eurofins Genomics Europe Sequencing GmbH (Munich, Germany).

A gene encoding SpyCatcher was amplified by PCR using the oligonucleotides, FW_BamHI_SpyC and RV_XhoI_SpyC, as primers and pET28_H-SpyCat as a template. The gene was subcloned into pACYC_Ptet_H-SUMO-mCherry-TRAP-K35C[10] via BamHI and XhoI sites, yielding pACYC_Ptet_H-SUMO-SpyC-TRAP-K35C. The N- and C-terminal segments of the TRAP-K35C gene as well as the SpyCatcher gene were individually amplified in the first PCR, using pACYC_Ptet_H-SUMO-SpyC-TRAP-K35C as a template, and the FW_XhoI_TRAP-N and RV_SacI_TRAP-N oligonucleotides as primers for the N-terminal segment, the FW_KpnI_TRAP-C and RV_MluI_TRAP oligonucleotides as primers for the C-terminal segment, and FW_SacI_SpyC and RV_KpnI_SpyC oligonucleotides as primers for the SpyCatcher gene. The second PCR was performed using equal amounts of the first PCR products as templates and the FW_BamHI_GGS and RV_MluI_TRAP oligonucleotides as primers. The second PCR product was then cloned into pACYC_Ptet_H-SUMO-SpyC-TRAP-K35C via the MluI and BamHI sites, yielding pACTet_H-SUMO-TRAP-K35C-loopSpyC.

The monomeric superfolder GFP, msfGFP, gene was prepared by introducing the V206K mutation on pACYC_H-sfGFP[23] using QuickChange mutagenesis using oligonucleotides, FW_sfGFP_V206K and RV_sfGFP_V206K, as PCR primers, to give pACYC-H-msfGFP. The msfGFP gene was amplified by PCR using the oligonucleotides, FW_KpnI_sfGFP and RV_EcoRI_msfGFP, and pACYC_H-msfGFP as template, and subsequently subcloned into pBAD_HTSpot_GFP(-30), a bacterial expression vector for GFP(-30) possessing His-tag, TEV protease cleavage site, and spot-tag at the N-terminus, prepared from pBAD_H-tev-GFP(+19) and pACYC_H-GFP(-30),[17] to give pBAD_HTSpot-msfGFP. The HTSpot-msfGFP gene was amplified by PCR using the oligonucleotides, FW_NcoI_His and RV_pBAD, and subcloned into pET28a plasmid using NcoI and HindIII sites, yielding pET28_HTSpot-msfGFP. The gene of His-tag and TEV protease cleavage site was prepared by two-step PCR: in the first one, using the oligonucleotides,

FW_NcoI_His and RV_Spy_BamHI_tev, as primers and pET28_HTSpot-msfGFP as template; the second using FW_NcoI_His and RV_KpnI_Spy2 as primers and the first PCR product as template. The second PCR product was subsequently subcloned into pET28_HTSpot-msfGFP, yielding pET28_H-SpyT-msfGFP.

The SrtA gene was amplified by PCR using the oligonucleotides FW_KpnI_SrtA and RV_KpnI_SrtA, as well as pET30b(+)_H-SrtA as template. The PCR product was subsequently subcloned into pET28_H-SpyT-msfGFP vector via KpnI and EcoRI sites, yielding pET28_H-SpyT-SrtA.

***Table S1***. Oligonucleotides used in this study

| Name | Sequence |
|---|---|
| FW_BamHI_SpyC | **GAGGGATCCGATAGCGCAACCCACATC** |
| RV_XhoI_SpyC | **GTAGCTCGAGCCAATATGCGCATCGCCTTTGG** |
| FW_XhoI_TRAP-N | **GGCTCGAGCTACACCAACTCTGACTTCGTTG** |
| RV_SacI_TRAP-N | **CTATCCGAGCTCCCGGTGAACTGCGCGATCAG** |
| FW_KpnI_TRAP-C | **CCGGGTACCGAACACACTAGTGCGATCAAAG** |
| RV_MluI_TRAP | **CTCACGCGTTATTTTTTACCTTCAGATTCGATAACAC** |
| FW_SacI_SpyC | **CACCGGGAGCTCGGATAGCGCAACCCACATCAAATTC** |
| RV_KpnI_SpyC | **CTAGTGTGTTCGGTACCCGGAATATGCGC** |
| FW_BamHI_GGS | **GAGGGATCCGGCTCAGGCGGCTCGAGCTACACCAAC** |
| FW_sfGFP-V206K | **AACCATTACCTGTCGACACAATCTAAGCTTTCGAAAGATCCCAACGAAAAGC** |
| RV_sfGFP-V206K | **GCTTTTCGTTGGGATCTTTCGAAAGCTTAGATTGTGTCGACAGGTAATGGTTG** |
| FW_KpnI_sfGFP | **CGGGTACCGCTAGCAAAGGAGAAGAACTTTTCAC** |
| RV_EcoRI_msfGFP | **GCTCGAATTCATTATTTGTAGAGTTCATCCATGCCAT** |
| FW_NcoI_His | **GAGCCATGGGCAGCAGCCATCATCATCATCATCATGGTATG** |
| RV_pBAD | **CTGATTTAATCTGTATCAGGCTG** |
| RV_Spy_BamHI_tev | **GGTCGGTTTGTACGCGTCAACCATAACGATGTGCGCGGATCCCTGGAAGTACA GGTTC** |
| RV_KpnI_Spy2 | **GCGGTACCCGAACCTTTGGTCGGTTTGTACGCGTCAACC** |
| FW_KpnI_SrtA | **GGAGGTACCATGCAAGCTAAACCTCAAATTCCGAAAG** |
| RV_KpnI_SrtA | **GGAATTGAATTCTCACTCGAGTTTGACTTCTGTAGCTACAAAGA** |

**Table S2. Plasmids used for protein production in this study**

| Name | Gene of interest | Tag | | Promoter/Operator[a] | Marker[b] |
|---|---|---|---|---|---|
| | | N- | C- | | |
| pET21_TRAP-K35C | TRAP-K35C | - | - | $P_{T7}$ / *lac*O | $Amp^R$ |
| pACYC_Ptet_H-SUMO-SpyC-TRAP-K35C | SpyCatcher-TRAP-K35C | His6-SUMO | - | $P_{tet}$ / *tet*O | $Cm^R$ |
| pACYC_Ptet_H-SUMO-TRAP-K35C (loopSpyC) | TRAP-K35C -loopSpyCatcher | His6-SUMO | - | $P_{tet}$ / *tet*O | $Cm^R$ |
| pET28_H-SpyT-msfGFP | msfGFP | His6-SpyTag | - | $P_{T7}$ / *lac*O | $Kan^R$ |
| pMAL_H-SpyT-sTEVp-R5 | sTEVp | His6-SpyTag | Arg5 | $P_{lac}$ / *lac*O | $Kan^R$ |
| pET28_H-SpyT-SrtA | SrtA | His6-SpyTag | - | $P_{T7}$ / *lac*O | $Kan^R$ |
| pET28_H-SpyT-hCAII | hCAII | His6-SpyTag | - | $P_{T7}$ / *lac*O | $Kan^R$ |

[a] $P_{T7}$/*lacO*, T7 promoter combined with lactose operator; $P_{tet}$/*tetO*, the tetracycline promoter combined with tetracycline operator; $P_{lac}$ / *lacO* , the lac promoter combined with lactose operator.

[b] $Amp^R$, ampicillin resistance; $Cm^R$, chloramphenicol resistance; $Kan^R$, kanamycin resistance.

**Table S3. Proteins used in this study**

Light blue = SpyCatcher

Green = msfGFP

Blue = SpyTag

Purple = Enzymes

**TRAP-K35C**

MYTNSDFVVIKALEDGVNVIGLTRGADTRFHHSECLDKGEVLIAQFTEHTSAIKVRGKAYIQTRH
GVIESEGKK*

**H-SUMO-SpyC-TRAP-K35C**

MHHHHHHGSSMASMKDHLIHNHHKHEHAHAEHLGSDSEVNQEAKPEVKPEVKPETHINLKVSD
GSSEIFFKIKKTTPLRRLMEAFAKRQGKEMDSLRFLYDGIRIQADQTPEDLDMEDNDIIEAHREQI
GGSDSATHIKFSKRDEDGKELAGATMELRDSSGKTISTWISDGQVKDFYLYPGKYTFVETAAPD
GYEVATAITFTVNEQGQVTVNGKATKGDAHIGSSYTNSDFVVIKALEDGVNVIGLTRGADTRFH
HSECLDKGEVLIAQFTEHTSAIKVRGKAYIQTRHGVIESEGKK*

**SpyC-TRAP-K35C**

SDSATHIKFSKRDEDGKELAGATMELRDSSGKTISTWISDGQVKDFYLYPGKYTFVETAAPDGYE
VATAITFTVNEQGQVTVNGKATKGDAHIGSSYTNSDFVVIKALEDGVNVIGLTRGADTRFHHSE
CLDKGEVLIAQFTEHTSAIKVRGKAYIQTRHGVIESEGKK*

**H-SUMO-TRAP-K35C-loopSpyC**

MHHHHHHGSSMASMKDHLIHNHHKHEHAHAEHLGSDSEVNQEAKPEVKPEVKPETHINLKVSD
GSSEIFFKIKKTTPLRRLMEAFAKRQGKEMDSLRFLYDGIRIQADQTPEDLDMEDNDIIEAHREQI
GGSGSGGSSYTNSDFVVIKALEDGVNVIGLTRGADTRFHHSECLDKGEVLIAQFTGSSDSATHIKF
SKRDEDGKELAGATMELRDSSGKTISTWISDGQVKDFYLYPGKYTFVETAAPDGYEVATAITFT
VNEQGQVTVNGKATKGDAHIPGTEHTSAIKVRGKAYIQTRHGVIESEGKK*

**TRAP-K35C-loopSpyC**

SGSGGSSYTNSDFVVIKALEDGVNVIGLTRGADTRFHHSECLDKGEVLIAQFTGSSDSATHIKFSK
RDEDGKELAGATMELRDSSGKTISTWISDGQVKDFYLYPGKYTFVETAAPDGYEVATAITFTVN
EQGQVTVNGKATKGDAHIPGTEHTSAIKVRGKAYIQTRHGVIESEGKK*

**H-SpyT-msfGFP**

MGSSHHHHHHGGSAHIVMVDAYKPTKGSGTASKGEELFTGVVPILVELDGDVNGHKFSVRGEG
EGDATNGKLTLKFICTTGKLPVPWPTLVTTLTYGVQCFSRYPDHMKRHDFFKSAMPEGYVQERT
ISFKDDGTYKTRAEVKFEGDTLVNRIELKGIDFKEDGNILGHKLEYNFNSHNVYITADKQKNGIK
ANFKIRHNVEDGSVQLADHYQQNTPIGDGPVLLPDNHYLSTQSKLSKDPNEKRDHMVLLEFVTA
AGITHGMDELYK*

**H-SpyT-sTEVp-R5**

GHHHHHHHGGSAHIVMVDAYKPTKGSGTAGESLFKGPRDYNPISSTIVHLTNESDGHTTSLYGIG
FGPFIITNKHLFRRNNGTLVVQSLHGVFKVKNTTTLQQHLIDGRDMIIIRMPKDFPPFPQKLKFREP
QREERIVLVTTNFQTKSMSSMVSDTSSTFPSGDGIFWKHWIQTKDGQCGSPLVSTRDGFIVGIHSA
SNFTNTNNYFTSVPKNFMELLTNQEAQQWVSGWRLNADSVLWGGHKVFMDKPEEPFQPVKEA
TQLMNRRRRR*

**H-SpyT-SrtA***

MGSSHHHHHHGGSAHIVMVDAYKPTKGSGTMQAKPQIPKDKSKVAGYIEIPDADIKEPVYPGPA
TREQLNRGVSFAKENQSLDDQNISIAGHTFIDRPNYQFTNLKAAKKGSMVYFKVGNETRKYKMT
SIRNVKPTAVEVLDEQKGKDKQLTLITCDDYNEETGVWETRKIFVATEVKLE*

**H-SpyT-hCAII**

MGSSHHHHHHGGSAHIVMVDAYKPTKGSGTAHHWGYGKHNGPEHWHKDFPIAKGERQSPVDI
DTHTAKYDPSLKPLSVSYDQATSLRILNNGHTFNVEFDDSQDKAVLKGGPLDGTYRLIQFHFHW
GSHDGQGSEHTVDKKKYAAELHLVHWNTKYGDFGKAVQQPDGLAVLGIFLKVGSANPGLQKV
VDVLDSIKTKGKSADFTNFDPRGLLPESLDYWTYPGSLTTPPLLECVTWIVLKEPISVSSEQVSKF
RKLNFNGEGEPEEPMVDNWRPTQPLKNRQIKASFK*

**Protein production and purification.**

TRAP 11mer patchwork assemblies containing SpyCatcher moieties were produced in *E. coli* strain BL21-Star(DE3) (Thermo Fisher Scientific) transformed with pET21_TRAP-K35C and pACYC_Ptet_H-SUMO-SpyC-TRAP-K35C or pACYC_Ptet_H-SUMO-TRAP-K35C (loopSpyC). Cells were cultured at 37 °C in lysogeny broth, Miller formulation (LB) medium supplemented with 100 µg/mL ampicillin and 25 µg/mL chloramphenicol until the OD600 reached ~0.7, at which point protein production was induced by the addition of IPTG (0.2 mM) and tetracycline (30 ng/mL). After culturing at 25 °C for ~18 h, cells were harvested by centrifugation at 5,000 ×g and 4 °C for 10 min, and stored at -20 °C.

For purification, the pellet from 400-mL culture was resuspended in 35 mL lysis buffer (50 mM sodium phosphate buffer (pH 8.0) containing 600 mM NaCl) supplemented with lysozyme (0.1 mg/mL), DNaseI (5 µg/mL), RNaseA (5 µg/mL), and protease-inhibitor-cocktail. After 1h at room temperature, the cells were lysed by sonication on a VCX-750 ultrasonic processor (Sonics and Materials, Newtown, CT, USA) equipped with a dual probe using a 50% duty cycle (2 sec pulse and 2 sec pose) and a 40% amplitude setting for 20 min on ice. After removal of the insoluble fraction by centrifugation at 16,600 ×g and 25 °C, the supernatant was loaded onto 2 mL of Ni(II)-NTA agarose resin in a gravity flow column. After washing with lysis buffer and wash buffer (50 mM sodium phosphate buffer (pH 8.0) containing 550 mM NaCl and 40 mM imidazole), the protein was eluted with 50 mM sodium phosphate buffer (pH 8.0) containing 300 mM NaCl and 250 mM Imidazole. Excess imidazole was removed by ultrafiltration (Amicon-15, 50K MWCO; Merck Millipore) using 50 mM sodium phosphate buffer (pH 8.0) containing 200 mM NaCl. The isolated protein was supplemented with 2 mM dithiothreitol (DTT) and treated with SUMO protease I (1 µg per 1 mg of target protein) at 4 °C overnight. Ni(II)-NTA agarose resin (100 µL) was added to the reaction and the supernatant was buffer exchanged to 50 mM sodium phosphate buffer (pH 8.0) containing 200 mM NaCl by ultrafiltration (Amicon-0.5, 30K MWCO). The resulting protein sample was further purified by size-exclusion chromatography on an NGC liquid chromatography pump system (Biorad) at room temperature [column, Superdex 200 increase 10/300 (Cytiva, Washington D.C., USA); eluent, 50 mM sodium phosphate buffer (pH 8.0) containing 200 mM NaCl; flow rate, 0.75 mL/min; room temperature; detection, absorbance at 280]. The fractions for the major peak were pooled and treated with 10 mM DTT, 1 mM L-tryptophan, and 2 mM EDTA, followed by buffer exchanged to 50 mM sodium phosphate buffer (pH 8.0) containing 200 mM NaCl by ultrafiltration (Amicon-4, 50K MWCO). Protein concentration was determined from the absorbance at 280 nm, $\varepsilon_{280}$ = 8,250 $M^{-1}$ $cm^{-1}$ or 19,060 $M^{-1}$ $cm^{-1}$ for TRAP as Trp-bound form or its fusion with SpyCatcher, respectively (http://pepcalc.com/). The stoichiometry of the patchwork assembly was estimated by the SDS-PAGE band densities analyzed on Image J software, resulting in 1.4 ± 0.3 and 1.1 ± 0.1 SpyCatcher per TRAP 11mer for the N-terminal and loop insertion variants, respectively.

N-terminally His- and Spy-tagged msfGFP and enzymes were produced and purified using an analogous protocol for TRAP, but without the His-SUMO-tag cleavage process. For enzyme production, cells were cultured at 18 °C for 48 h. After purification by size-exclusion chromatography, proteins were buffer-exchanged to phosphate-buffered saline (PBS), added with glycerol to a final concentration of 20% (v/v), flash-frozen with liquid nitrogen, and stored at -80 °C. GFP concentration was determined by absorbance at 488 nm using the extinction coefficient, $\varepsilon_{488}$ = 54,900 ± 2,100 $M^{-1}$ $cm^{-1}$, which was estimated based on the absorbance of the chromophore obtained following protein denaturation in 0.1 M NaOH ($\varepsilon_{447,den}$ = 44,100 $M^{-1}$ $cm^{-1}$).[24] Enzyme concentration was estimated by absorbance at 280 nm using the theoretical extinction coefficients (sTEVp, 33,570 $M^{-1}$ $cm^{-1}$; SrtA, 14,650 $M^{-1}$ $cm^{-1}$; hCAII, 51,350 $M^{-1}$ $cm^{-1}$).

**TRAP-cage assembly**

Au(I)-mediated cage assembly was performed by mixing TRAP (500 µM) with TPPMS-Au(I)-Cl (500 µM, 1 equiv.) in 50 mM sodium phosphate buffer (pH 8.0) containing 600 mM NaCl for room temperature overnight. For DTME-mediated assembly, TRAP (200 µM) was added with DTME (1 mM, 5 equiv.). The reaction was performed in 50 mM sodium phosphate buffer (pH 7.4) containing 2.5% (v/v) DMF at room temperature for 1 h, followed by treatment with $Na_2CO_3$ (25 mM), final pH ~10, at room temperature for 30 min to hydrolyze maleimide moieties. For both cases, after buffer-exchange to PBS by ultrafiltration (Amicon-4, 50K MWCO), assembled TRAP-cages were isolated by size-exclusion chromatography [column, Superose 6 increase 10/300 GL (Cytiva); eluent, PBS; flow rate, 0.75 mL/min; room temperature; detection, absorbance at 280].

**GFP encapsulation in TRAP$^{Au(I)}$-cage**

Purified H-SpyT-msfGFP (20 µM) were mixed with TRAP-cage containing SpyCatcher moiety (20 µM, respect to SpyCatcher concentration) in PBS at room temperature overnight, unless specified. Assembled TRAP-cage was isolated by size-exclusion chromatography [column, Superose 6 increase 10/300 GL; eluent, PBS; flow rate, 0.75 mL/min; room temperature; detection, absorbance at 280 and 488 nm], and analyzed by negative-stain transmission electron microscopy (TEM), native polyacrylamide gel electrophoresis (Native-PAGE), SDS-PAGE, and absorbance spectra. Encapsulation experiments were performed with 4 different batches of TRAP, and mean ± standard deviation is provided for quantitative assays.

For guest titration experiments, the different concentrations of H-SpyT-msfGFP (0, 5, 10, 20, or 30 µM) were mixed with TRAP-cage containing the SpyCatcher moiety (20 µM, with respect to SpyCatcher concentration) in PBS at room temperature, overnight. For kinetics, encapsulation was performed at 4 °C or room temperature at 0, 1, 2, 4, 20 h. The resulting mixtures were directly analyzed by Native- and SDS-PAGE. The titration and kinetics experiments were performed with two different batches of TRAP, and mean ± standard deviation is provided.

**Negative-stain transmission electron microscopy (TEM)**

Protein samples in PBS (0.035 mg/mL, with respect to TRAP concentration) were placed on a glow-discharge-treated carbon-coated copper grid (EM resolution, Sheffield, UK). Excess solution was removed using filter paper followed by incubation with 2% (w/v) uranyl acetate. Samples were visualized on a JEOL JEM-1230 electron microscope with 80 kV operation. Obtained images were analyzed using Image J software.

**Native-PAGE**

Samples were prepared in a native-PAGE loading buffer (50 mM BisTris-HCl buffer (pH 7.2) containing 10% (v/v) glycerol, 0.004% (w/v) bromophenol blue). Electrophoresis was performed using 3–12% native Bis-Tris gels and NativePAGE running buffer (both from Thermo Fisher Scientific) at 150 V for 90 min. Fluorescent bands were visualized using a ChemiDoc MP (Bio-Rad) in fluorescein detection mode (460–490 nm excitation and 532/28 nm emission filters). The same gels were also stained with ReadyBlue Protein Gel Stain

(Sigma-Aldrich). For analysis of the samples without GFP, 0.08% (w/v) Coomassie Blue was added to the loading and running buffer (Blue-Native-PAGE).[25]

**SDS-PAGE**

Samples were prepared in SDS-PAGE sample buffer (62.5 mm Tris-HCl buffer (pH 6.8) containing 2.5% (w/v) sodium dodecyl sulfate (SDS), 10% (v/v) glycerol, 5% (v/v) β-mercaptoethanol, 0.2 mM DTT, and 0.004% (w/v) bromophenol blue) with 5 M urea addition and heated at 95 ºC for 20 min. Potential carbamylation due to the presence of urea was taken into account, but it did not lead to a significant resolution decrease. Electrophoresis was performed using 15% polyacrylamide gels and a Tris-Tricine buffer system[26] at 80 V for 10 min followed by 150 V for 60 min. As a protein standard, PageRuler™ Prestained Protein Ladder (Thermo Fisher Scientific) was used. Protein bands were visualized using ReadyBlue Protein Gel Stain (Sigma-Aldrich). For densitometry analysis, the images were analyzed using Image J software.

**GFP encapsulation efficiency**

The average number of GFP encapsulated in a TRAP-cage was estimated based on the absorbance ratio at 280/488 nm as previously reported.[5d] The spectra were recorded on a Shimadzu UV1900 UV-Vis spectrophotometer (Kyoto, Japan) using a 1-cm-light-pass-quartz cuvette at 25 °C. The extinction coefficient of the H-SpyT-msfGFP was determined using the observed absorbance ratio at 280/488 nm to be $\varepsilon_{G280}$ = 24,000 ± 1,900 $M^{-1}$ $cm^{-1}$.

**Enzyme encapsulation**

All the enzymes were treated with DTT (10 mM) for 1 h, followed by buffer exchange to PBS by ultrafiltration (Amicon-4, 50K MWCO) prior to the encapsulation study. Additional $ZnSO_4$ (1 equiv.) treatment was also conducted for hCAII. An enzyme (8 µM or 16 µM for sTEVp and hCAII or SrtA, respectively) was mixed with TRAP containing SpyCatcher (32 µM respect to SpyCatcher concentration, 4 or 2 equiv.) in either 11mer or assembled cage format in PBS (for hCAII, in 25 mM HEPES-KOH buffer pH 7.5 containing 100 mM NaCl) at 4 °C overnight. The resulting mixture was analyzed by Native- and SDS-PAGE, negative-stain TEM, and directly used for the catalytic activity test without further purification. For sTEVp, the sample after encapsulation was treated with 10 mM DTT another overnight to test the catalytic activity upon cage disassembly. Encapsulation experiments were performed with 2 different batches of TRAP, and mean ± standard deviation is provided for quantitative assays.

**Enzyme kinetics**

The activity of the cysteine-proteases, sTEVp and SrtA, was measured using FRET substrates containing aminobenzoic acid (Abz) and 2,4-dinitrophenol (Dnp) as previously reported.[17] Peptide concentrations were determined by absorption at 365 nm ($\varepsilon_{365}$ = 17,300 $M^{-1}cm^{-1}$).[27] 3.75-120 µM substrate solution in 50 mM sodium phosphate buffer (pH 7.4) containing 1 mM EDTA and 2% (v/v) DMF was added with enzyme stock solution (final 40 nM or 200 nM for sTEVp or SrtA, respectively), and then time-dependent fluorescence was monitored at 25 °C with excitation at 320 nm and emission at 420 nm on a Shimadzu RF6000 fluorometer (Kyoto, Japan). Inner filter effects of Abz fluorescence were corrected using a method previously reported.[17] For hCAII, the esterase activity on 4-nitrophenyl acetate was measured. 0.16 -5 mM substrate solution in 25 mM HEPES-KOH buffer (pH 7.5) containing 100 mM NaCl and 5% (v/v) MeCN was added with enzyme stock solution (final 40 nM), and then time-dependent absorbance at 405 nm ($\varepsilon_{405}$ = 13,500 $M^{-1}$ $cm^{-1}$)[28] was monitored at 25 °C on a Shimadzu UV1900 UV-Vis spectrophotometer. For sTEVp, steady-state kinetic parameters, $k_{cat}$ and $K_M$, were determined by Michaelis-Menten curve fitting using GraphPad Prism. Because no saturation was observed for SrtA and hCAII under the tested conditions, the kinetic efficiency, $k_{cat}/K_M$, was determined by linear fitting using the data points no greater than 60 µM and 2.5 mM of substrate for SrtA and hCAII, respectively.
